# Interactions between cardiac activity and conscious somatosensory perception

**DOI:** 10.1101/529636

**Authors:** Paweł Motyka, Martin Grund, Norman Forschack, Esra Al, Arno Villringer, Michael Gaebler

**Author notes:** Corresponding author: Paweł Motyka, Faculty of Psychology, University of Warsaw, Stawki 5/7, 00-183 Warsaw, Poland.

## Abstract

Fluctuations in the heart’s activity can modulate the access of external stimuli to consciousness. The link between perceptual awareness and cardiac signals has been investigated mainly in the visual and auditory domain. We here investigated whether the phase of the cardiac cycle and the pre-stimulus heart rate influence conscious somatosensory perception. We also tested how conscious detection of somatosensory stimuli affects the heart rate. Electrocardiograms (ECG) of 33 healthy volunteers were recorded while applying near-threshold electrical pulses at a fixed intensity to the left index finger. Conscious detection was not uniformly distributed across the cardiac cycle but significantly higher in diastole than in systole. We found no evidence that the heart rate before a stimulus influenced its detection but hits (correctly detected somatosensory stimuli) led to a more pronounced cardiac deceleration than misses. Our findings demonstrate interactions between cardiac activity and conscious somatosensory perception, which highlights the importance of internal bodily states for sensory processing beyond the auditory and visual domain.

**Impact Statement:** It is highly debated to what extent cardiac activity modulates the access of external stimuli to consciousness. The evidence is inconsistent across sensory modalities and previous research focused at specific intervals within the cardiac cycle. Here, we examined the perception of near-threshold electrical pulses across the entire cardiac cycle. Our results show that conscious somatosensory perception is enhanced during the late phase of the cardiac cycle (at diastole) and associated with a more pronounced cardiac deceleration (as compared to non-detected stimuli). This strengthens the evidence that the physiological state of the body influences how we perceive the world.

## 1. INTRODUCTION

The internal state of the body is continuously monitored by interoceptive regions and networks in the brain (Craig, 2009; Barrett & Simmons, 2015; Kleckner et al., 2017). Besides their well-described role in homeostatic regulation, visceral signals have been argued to contribute to a wide range of psychological phenomena, including emotions (Critchley & Garfinkel, 2017; Wiens, 2005), empathy (Grynberg & Pollatos, 2015; Fukushima, Terasawa, & Umeda, 2011), time perception (Di Lernia et al., 2018; Meissner & Wittmann, 2011), self-consciousness (Craig, 2009; Park & Tallon-Baudry, 2014), and decision-making (Gu & Fitzgerald, 2014; Seth, 2014). At the perceptual level, it remains unclear to what extent interoceptive states can modulate the conscious access to sensory signals. Here, we examined the interactions between perceptual awareness for somatosensory stimuli and cardiac activity – that is, the phase of the cardiac cycle and the heart rate.

The cardiac cycle from one heartbeat to the next can be divided into two phases: systole, when the heart contracts and ejects blood into the arteries – leading to activation of pressure-sensitive baroreceptors in arterial vessel walls – and diastole, when the cardiac muscle relaxes, the heart refills with blood, and baroreceptors remain quiescent (Landgren, 1952; Mancia & Mark, 2011). Baroreceptor activity signals the strength and timing of each heartbeat to the nuclei in the lower brain stem, where the signal is relayed to subcortical and cortical brain regions (Dampney, 2016). In studies with non-invasive baroreceptor stimulation, their activity was found to decrease BOLD signal (Makovac et al., 2015) and ERP amplitudes (Rau, Pauli, Brody, Elbert, & Birbaumer, 1993; Rau & Elbert, 2001) in cortical regions. Baroreceptor firing is thought to underlie cardiac cycle effects on behavior and cognition (Duschek, Werner, & Reyes Del Paso, 2013; Garfinkel & Critchley, 2016), like decreased intensity ratings for acoustic (Cohen, Lieb, & Riest, 1980; Schulz et al., 2009) or painful stimulation (Wilkinson et al., 2013) as well as higher reaction times to stimuli (Birren, Cardon, & Philips, 1963; Edwards, Ring, McIntyre, Carroll, & Martin, 2007; McIntyre, Ring, Edwards, & Carroll, 2008) during early (i.e., at systole) compared to later phases (i.e., at diastole) of the cardiac cycle.

There are conflicting findings of whether the cardiac cycle modulates the access of exteroceptive information to perceptual awareness. Earlier studies reported that the detection of visual (Réquin & Brouchon, 1964, Sandman, McCanne, Kaiser, & Diamond, 1977) and auditory signals (Saxon, 1970) vary for different points of the cardiac cycle. However, other studies in the visual (Elliot & Graf, 1972) and auditory domain (Delfini & Campos, 1972; Velden & Juris, 1975) did not find such variations. More recently, an enhanced detection selectively for fearful faces was observed during cardiac systole (Garfinkel et al., 2014). As almost all studies in that field involved visual or auditory stimuli, it remains unclear whether cardiac-phase related fluctuations occur in other sensory modalities. The only previous study in the somatosensory domain with a behavioral measure of perception reported lower detection thresholds for electrical stimulation at systole compared to diastole (Edwards, Ring, McIntyre, Winer, & Martin, 2009). As in most studies of cardiac-phase effects, the detection performance was sampled only at fixed time points (R+0, R+300, and R+600 ms), which may have missed perceptual changes at other parts of the cardiac cycle.

In the present study, we examined fluctuations in conscious somatosensory perception across the entire cardiac cycle. Given the variations in cortical excitability over the cardiac cycle, we hypothesized that detection of near-threshold electrical stimuli is not equally distributed but varies over the interval between one heartbeat and the next. We also aimed to explore associations between conscious somatosensory perception and the heart rate. The bidirectional information flow between the heart and the brain (Faes et al., 2017, Lin, Liu, Bartsch, & Ivanov, 2016, Valenza, Toschi, & Barbieri, 2016) implies that cardiac activity may not only impact perception but also be influenced by it. Therefore, we tested whether the pre-stimulus heart rate influences conscious perception and, in turn, whether perception changes the (post-stimulus) heart rate.

Regarding the relation between the heart rate and perception, an early theory suggested that a decreased heart rate increases sensitivity to sensory stimulation by directing attention to external rather than internal signals (Graham & Clifton, 1965; Lacey, Kagan, Lacey, & Moss, 1963; Lacey, 1967; Sandman, 1986). The evidence for this hypothesis is mixed and comes only from studies in the auditory and visual domain: For auditory thresholds, there were no differences between transient periods of low and high heart rate (Edwards & Alsip, 1969) unless the procedure involved exercise-induced changes in heart rate (Saxon & Dahle, 1971). In addition to such heart rate variations over longer periods of time, quick changes from one heartbeat to the next were suggested to modulate perception (Lacey & Lacey, 1974; Sandman et al., 1977). In general, cardiac deceleration (i.e., a lengthening of the period between consecutive heartbeats) is known to occur in anticipation of a (cued) stimulus or in reaction to a salient stimulus (Lacey & Lacey, 1970, 1977; Simons, 1988), and it is typically followed by cardiac acceleration after the behavioral response (e.g., Börger & Meere, 2000; Park, Correia, Ducorps, & Tallon-Baudry, 2014). While both spontaneous (Sandman et al., 1977) and conditioned (McCanne & Sandman, 1974) cardiac deceleration coincident with a visual stimulus was found to increase its detection, other – more recent – studies did not show a modulation of visual awareness by heart rate changes prior to and coincident with a near-threshold stimulus (Cobos, Guerra, Vila, & Chica, 2018; Park et al., 2014).

For heart rate changes *after* stimulus presentation, earlier studies found a cardiac deceleration in response to suprathreshold visual (Davis & Buchwald, 1957), auditory (Davis, Buchwald, & Frankmann, 1955; Uno & Grings, 1965; Wilson, 1964), tactile (Davis et al., 1955), and olfactory stimuli (Gray & Crowell, 1968). Additionally, cardiac deceleration was found to be more pronounced after viewing unpleasant compared to pleasant or neutral scenes (Bradley, Cuthbert, & Lang, 1990; Greenwald, Cook, & Lang; 1989; Hare, 1973; Libby, Lacey, & Lacey, 1973; Walker & Sandman, 1977). Most importantly in the context of this work, recent studies using near-threshold visual stimuli showed that hits resulted in increased cardiac deceleration compared to misses (Cobos et al., 2018; Park et al., 2014). This suggests that not only the physical characteristics of a stimulus determine the cardiac response but also the level of its processing (i.e., conscious vs. nonconscious).

The association between cardiac activity and perception was also related to cardiac-phase independent variations in arterial pressure after changes in heart rate (Sandman et al., 1977). In this view, the late phase of the cardiac cycle (i.e., diastole) and cardiac deceleration result in – similar but not identical – transient decreases in blood pressure; thus facilitating the access of external stimuli to consciousness by decreasing the inhibitory effects of baroreceptor activity on the brain (Sandman, 1986; Sandman et al., 1977). Notably, even though higher mean arterial blood pressure has been associated with higher resting heart rate (Christofaro, Casonatto, Vanderlei, Cucato, & Dias, 2017; Mancia et al., 1983), increases in blood pressure after cardiac deceleration (i.e., decreases in heart rate) were observed during experimental tasks (Otten, Gaillard, & Wientjes, 1995; Wölk, Velden, Zimmermann, & Krug. 1989). In addition, animal studies showed that – also with constant mean arterial pressure – the heart rate elevation leads to an increased discharge of arterial baroreceptors (Abboud & Chapleau, 1988; Barrett & Bolter, 2006). Taken together, these findings suggest that the heart rate contributes to cortical excitability through a transient modulation of baroreceptor activity.

Furthermore, we aimed to test whether the influence of cardiac signals on perception varies with inter-individual differences in interoceptive accuracy, that is, the ability to consciously perceive signals originating from the body (Garfinkel, Seth, Barrett, Suzuki, & Critchley, 2015). Given that the capacity to detect one’s own heartbeat has been repeatedly shown to modulate (usually strengthen) cardiac effects on perception and behavior (Critchley & Garfinkel, 2018; Dunn et al., 2010; Suzuki, Garfinkel, Critchley, & Seth, 2013), we hypothesized that the link between conscious somatosensory perception and cardiac activity would be stronger for participants with higher interoceptive accuracy (measured with the Heartbeat Counting Task; Schandry, 1981).

In sum, given that baroreceptor activity, which is thought to suppress the processing of external input, varies both across the cardiac cycle and with the heart rate, we hypothesized that perceptual awareness for somatosensory stimuli fluctuates as a function of these cardiac parameters. Also, we explored whether a consciously detected somatosensory stimulus affects the heart rate differently compared to a non-detected stimulus and whether cardiac effects on conscious somatosensory perception varies with the capacity to consciously perceive one’s heartbeat.

## 2. METHOD

### 2.1 Participants

Thirty-three healthy volunteers (17 females, mean age = 25.9, *SD* = 4.1, range: 19-36 years, right-handed) were recruited from the database of the Max Planck Institute for Human Cognitive and Brain Sciences in Leipzig, Germany. The procedure was approved by the ethics committee of the Medical Faculty at the University of Leipzig. All participants gave written informed consent before taking part in the study and were financially compensated for their participation.

### 2.2 Apparatus

Electrocardiography (ECG) was measured while near-threshold electrical finger nerve stimulation was applied. ECG was recorded at a sampling frequency of 1000 Hz with a Brain Products BrainAmp (Brain Products GmbH, Gilching, Germany). Electrodes were placed on the wrists and the left ankle (ground) according to Einthoven’s triangle. Electrical finger nerve stimulation was performed with a constant-current stimulator (DS5; Digitimer) applying single rectangular pulses with a length of 200 μs. A pair of steel wire ring electrodes was attached to the middle (anode) and the proximal (cathode) phalanx of the left index finger. The experiment was programmed and behavioral data was recorded with Matlab 8.5.1 (Psychtoolbox 3.0.11, Brainard, 1997; Pelli, 1997; Kleiner et al., 2007).

### 2.3 Procedure

Each participant was tested individually in a dimly lit experimental chamber seated in a comfortable chair and facing a computer screen. After a brief explanation of the experimental procedure and the attachment of ECG electrodes, the steel wire ring electrodes were attached to the left index finger. The response button box was placed under the right hand. The computer screen indicated when to expect a stimulus and when to respond (Fig. 1). Participants responded with “yes” if they felt an electrical stimulus and “no” if not. The left/right button-response mapping (yes-no or no-yes) was pseudo-randomized across participants. The experimental session consisted of 360 trials divided into three blocks. Each block included 100 trials with near-threshold stimulation and 20 catch trials without stimulation in pseudo-randomized order. The intensity of electrical stimulation was fixed throughout a block. Before each block, the somatosensory perceptual threshold was assessed using an automated staircase procedure to estimate a stimulus intensity that would be equally likely to be felt or not (the 50% detectability level). The applied method combines a coarser staircase procedure (“up/down method“) and a more fine-grained Bayesian procedure (“psi method“) of the Palamedes Toolbox (Kingdom and Prins, 2010). The automated threshold assessment resembled the actual experimental design, except for the shorter (500 ms) inter-trial interval and the time window in which stimulation could occur (1,000 ms). Thus, before each block, the experimenter made a data-driven decision of the individual sensory threshold (*M* = 2.24, *SD* = 0.81, range = 1-5 milliamperes). At the end of the experimental session, inter-individual differences in interoceptive accuracy (Garfinkel et al., 2015) were assessed with a Heartbeat Counting Task (Schandry, 1981), in which participants were asked to estimate the number of their heartbeats in five intervals of different duration (detailed in the Supplementary Material).

**Figure 1.**
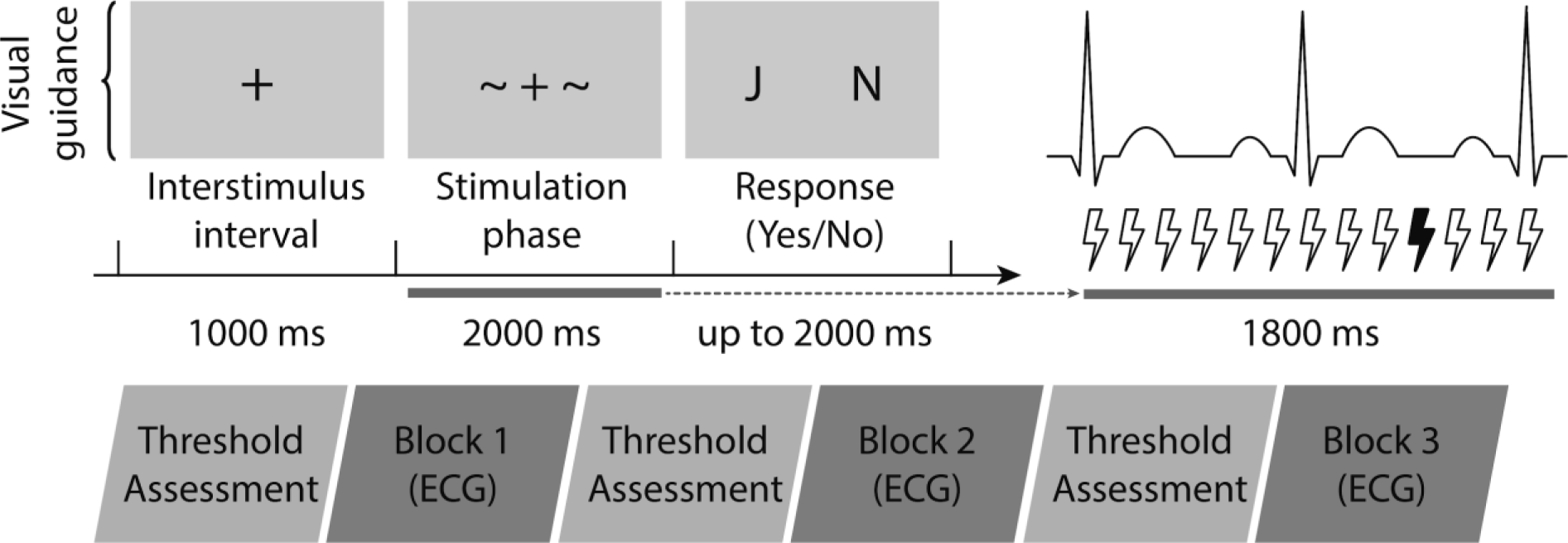
Near-threshold somatosensory signal detection task. Trial procedure (upper row): each trial started with a 1,000-ms central fixation cross, followed by the 2,000-ms time window during which the stimulus could occur (except for the first and the last 100 ms of this interval). The stimulation onset was pseudo-randomized within this 1,800-ms time window, aiming for a uniform distribution of stimuli over the entire cardiac cycle. Next, the response phase began (cued by displaying “J N” – corresponding to “yes” and “no”, respectively) and lasted until participants gave a response within the maximum time of 2,000 ms. After the button press, the fixation cross was visible for the rest of the 2,000-ms interval so that the total duration of each trial was kept constant at 5,000 ms. The next trial followed immediately so that the duration of each block was fixed (10 min). An experimental session (lower row) consisted of three such blocks, which were each preceded by a threshold assessment to estimate stimulus intensities with 50% detection probability.

### 2.4 Statistical analysis

All statistical analyses were conducted using R version 3.5.1 (R Core Team, 2016) with RStudio version 1.1.453 (RStudio Team, 2016) and the Circular package version 0.4.93 (Agostinelli & Lund, 2013). Kubios 2.2 (Tarvainen et al., 2014; Biosignal Analysis and Medical Imaging Group, Department of Applied Physics, University of Eastern Finland, Kuopio, http://kubios.uef.fi/) was used to automatically detect and visually inspect R peaks in the ECG. Falsely detected or missed R peaks (<0.2%) were manually corrected. A two-sided alpha level of 0.05 was used in all statistical analyses. All preprocessed data and the codes used for the main and supplementary analyses are available on GitHub at https://github.com/Pawel-Motyka/CCSomato.

#### 2.4.1 Behavior

Prior to the analysis, the following data was excluded: 191 trials (from 26 participants) with no response within two seconds (1.7% of all trials), 15 trials where the stimulation failed, 2 trials with the unassigned button pressed, and 2 trials with physiologically implausible IBI lengths (>1,500 ms). Also, one block of one participant was excluded due to data-recording failure. Thus, the total number of trials retained for analysis was 11,550 (from 33 participants): 4,530 hits (correctly detected near-threshold stimuli), 5,104 misses (not detected near-threshold stimuli), 81 false alarms (wrongly detected non-stimulation), and 1,835 correct rejections (correctly detected non-stimulation).

#### 2.4.2 ECG data

To investigate cardiac phase-related variations in perceptual awareness for somatosensory stimuli while accounting for both the oscillatory and the biphasic nature of cardiac activity, the distribution of hits and misses were examined (1) over the whole cardiac cycle by means of circular statistics (Pewsey, Neuhäuser, & Ruxton, 2013) and (2) by testing differences in hit rates between the two cardiac phases (systole and diastole), respectively. Furthermore, it was analyzed (3) whether pre- and post-stimulus changes in heart period differed between hits and misses.

1. Circular statistics allows to analyze the distribution of hits and misses along the entire cardiac cycle (from one R peak to the next). For each participant, the mean phase angle, at which hits or misses occurred on average, was calculated in degrees (see section “Determination of stimulus onset distribution across the cardiac cycle“). At the group level, it was tested with Rayleigh tests (Pewsey et al., 2013) whether the distributions of hits and misses deviated from the uniform distribution. The Rayleigh test is based on the mean vector length out of a sample of circular data points and specifies the average concentration of these phase values around the circle – ranging from 0 to 1 indicating no to perfect (angular) concentration, respectively. A statistically significant Rayleigh test result indicates that the data are unlikely to be uniformly distributed around the circle (in this case: the cardiac cycle).
2. Binary analysis, based on the segmentation of the cardiac cycle into the two cardiac phases allowed us to compare our results to previous studies of cardiac effects on perception. To divide the cardiac cycle into systole and diastole, the trial-specific cardiac phases were computed based on cardio-mechanical events related to the ECG signal (for a description of the applied t-wave end detection algorithm, see section 2.6 “Determination of individual cardiac phases“). Given the between-subject variation of cardiac phase lengths arising, for example, from differences in heart rate (Herzog et al., 2002; Lewis, Rittogers, Froester, & Boudoulas, 1977; Wallace, Mitchell, Skinner, & Sarnoff. 1963), an individualized approach was used – instead of rather arbitrary and fixed systole and diastole intervals (e.g., defining systole as the 300 ms following an R peak). Stimulus onsets were assigned to the corresponding cardiac phase (i.e., systole or diastole) for each trial. Then, for each participant, hit rates were calculated separately for systole and diastole. A paired *t* test was used to determine whether hit rates differed between cardiac phases.
3. To analyze the pre- and post-stimulus heart rate for hits and misses, the mean lengths of six consecutive interbeat intervals (IBIs) were computed (with an average IBI of 827 ms (*SD* = 119 ms), these aimed to cover the full trial length of 5,000 ms): two before the stimulation (*S-2*, *S-1*), one at which the stimulus occurred (*Stimulus*), and three after the stimulation (*S+1*, *S+2*, *S+3*). To test whether the (changes in) heart period differed between hits and misses, a two-way repeated-measures analysis of variance (ANOVA) was used – with perceptual awareness (hits/misses) and time (six IBIs, *S-2* to *S+3*, per trial) as factors – followed by post-hoc Bonferroni-corrected paired *t* tests. Furthermore, an association between the extent of cardiac deceleration and the conscious access to somatosensory stimuli was investigated. For each trial, cardiac deceleration was calculated (and z-scored within participants) as the difference between the lengths of the IBI at which the stimulus occurred (*Stimulus*) and the IBI prior to it (*S-1*). A paired *t* test was used to examine whether the extent of cardiac deceleration differed between hits and misses.

#### 2.4.3 Interoceptive accuracy

A score of interoceptive accuracy was calculated for each participant. The closer the estimated number to the number of heartbeats measured by the ECG over five intervals, the higher the interoceptive accuracy score (cf. Supplementary Material). The sample was then median-split into groups of high and low interoceptive accuracy, which were compared using analyses described in section 2.4.2.

### 2.5 Determination of stimulus onset distribution across the cardiac cycle

In each trial, stimulus onset was pseudo-randomized within a 1,800-ms time window. Stimulation at different points of IBI aimed to cover the entire cardiac cycle for each subject. For each stimulus, the time of the previous and the subsequent R peak were extracted from the ECG to calculate the stimulus onset’s relative position within the IBI using the following formula: [(*onset time – previous R peak time*) / (*subsequent R peak time* – *previous R peak time*)] × 360, assigning the values from 0 to 360 degrees (with 0 indicating the R peak before the stimulus). The distribution of stimulus onsets was tested separately for each participant with a Rayleigh test for uniformity. One participant was excluded from further circular analyses due to non-uniformly distributed stimulation onsets across the cardiac cycle, *R̅* = 0.11, *p* = 0.009. For the rest of the participants, the assumption of uniform onset distributions was fulfilled (all *p* > 0.091).

### 2.6 Determination of individual cardiac phases

To account for the biphasic nature of cardiac activity, we encoded the length of individual cardiac phases using the t-wave end detection method (Vázquez-Seisdedos, Neto, Marañón Reyes, Klautau, & Limão de Oliveira, 2011): First, the peak of the t-wave was located as a local maximum within a physiologically plausible interval (up to 350 ms after the R peak). Subsequently, a series of trapezes was calculated along the descending part of the t-wave signal, defining the point at which the trapezium’s area gets maximal as the t-wave end. Detection performance was visually controlled by overlaying the t-wave ends and the ECG trace from each trial. 27 trials with extreme systole lengths (more than 4 SDs above or below the participant-specific mean) were excluded.

Although mechanical systole cannot be fully equated with the duration of electrical systole in the ECG (Fridericia, 1920), both are closely tied under normal conditions (Boudoulas, Geleris, Lewis, & Rittgers, 1981; Coblentz, Harvey, Ferrer, Cournand, & Richards, 1949; Fridericia, 1920; Gill & Hoffman, 2010). Systolic contraction of the ventricles follows from their depolarisation (marked in the ECG by the QRS complex), whereas the closure of the aortic valve, terminating the systolic blood outflow, corresponds to ventricular repolarization (around the end of the t-wave; Gill & Hoffmann, 2010). In our study, the ventricular systolic phase (further referred to as “systole“) was defined as the time between the R peak of the QRS complex and the t-wave end, while diastole was defined as the remaining part of the RR interval.

## 3. RESULTS

### 3.1 Detection rate for near-threshold somatosensory stimuli

On average 46.7% of the near-threshold trials were detected (*SD* = 16.2%, range: 15.1-79.3%). The false alarm rate was 4.2% (*SD* = 5.7%, range: 0-16.6%).

### 3.2 Hits concentrated in the late phase of the cardiac cycle

Rayleigh tests were applied to analyze the distribution of hits and misses across the cardiac cycle. Hits were not uniformly distributed across the cardiac cycle, *R̅* = 0.32, *p* = 0.034 (Fig. 2A), with their mean angle directing to the later phase of the cardiac cycle (i.e., diastole). Misses showed a non-significant tendency to deviate from uniformity, *R̅* = 0.30, *p* = 0.060 (Fig. 2B), with their mean angle directing to the earlier phase of the cardiac cycle (i.e., systole). For 14 out of 32 participants, the individual mean angles for hits fell into the last quarter of the cardiac cycle. The individual mean angles for misses accumulated in the second quarter of the cardiac cycle for 13 participants. Distributions of hits or misses across the cardiac cycle did not differ significantly between participants with high or low interoceptive accuracy (see Fig. S1 in Supplementary Material).

**Figure 2.**
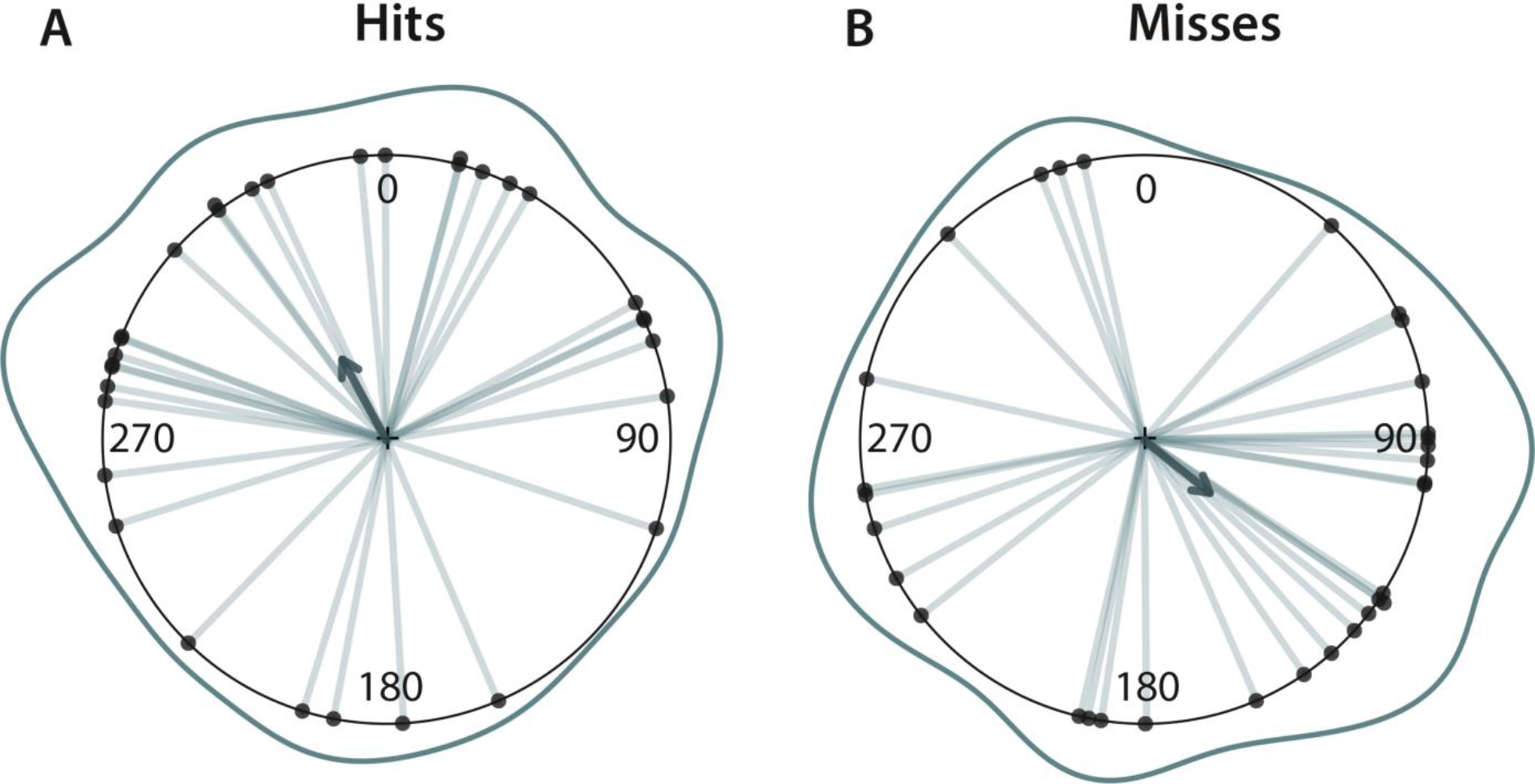
Distribution of (**A**) hits and (**B**) misses across the cardiac cycle (i.e., the interval between two R peaks; at 0/360°). Rayleigh tests showed a significant deviation from a uniform distribution for hits (*R̅* = 0.32, *p* = 0.034) and a non-significant trend for misses (*R̅* = 0.30, *p* = 0.060). Each dot (and line) indicates one participant’s mean phase angle. The annular line depicts the distribution of individual means. The darker arrows represent the directions of the group means for hits (331°) and misses (129°), with their length indicating the concentration of individual means across the cardiac cycle (hits: 0.32, misses: 0.30 – with 1 indicating perfect angular concentration).

### 3.3 Higher hit rates in diastole than in systole

Accounting for the biphasic nature of cardiac activity, differences in hit rates between systole and diastole were examined. Hit rates for near-threshold somatosensory stimuli were significantly higher during diastole (*M* = 47.9%, *SD* = 16.5%) than during systole (*M* = 45.1%, *SD* = 16.3%), *t*(32) = −2.76, *p* = 0.009, Cohen’s *d* = 0.48. Increased hit rate during diastole was observed for 25 out of 33 participants (Fig. 3). This mirrors the concentration of hits in the later phase of the cardiac cycle (see circular statistics, Fig. 2). Hit rates at different cardiac phases did not differ significantly between the groups with high and low interoceptive accuracy (see Fig. S2 in Supplementary Material).

**Figure 3.**
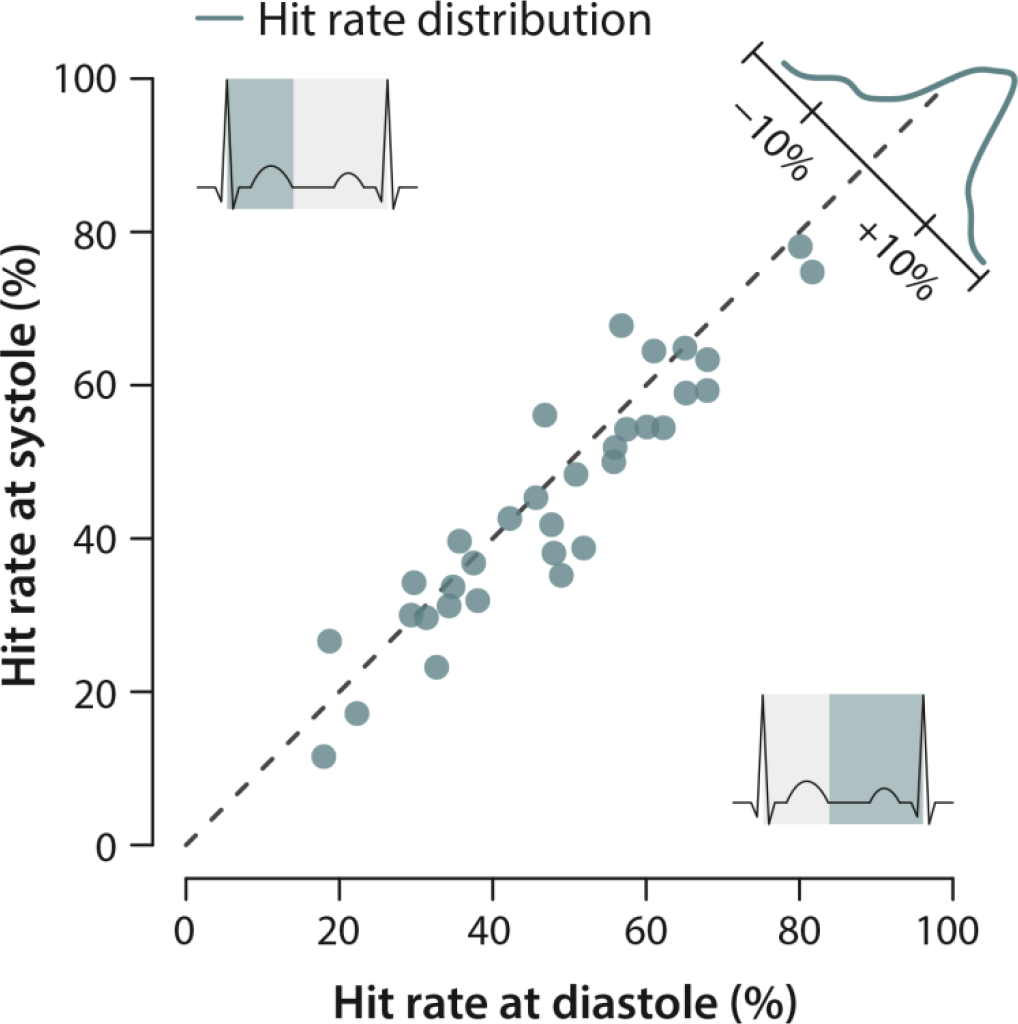
Higher hit rates in diastole than in systole. The coordinates of each dot represent a participant’s mean hit rate at systole (x-axis) and diastole (y-axis). The dashed lines mark the identity line in hit rate between cardiac phases. The distribution in the upper right corner aggregates the frequency across participants. The probability distribution was shifted towards diastole indicating significantly higher hit rates during the later phase (i.e., diastole) than the earlier phase (i.e., systole) of the cardiac cycle (*t*(32) = −2.76, *p* = 0.009, Cohen’s *d* = 0.48).

### 3.4 Pre- and post-stimulus heart rate (changes) for hits and misses

To investigate how the heart interacts with conscious somatosensory perception, pre- and post-stimulus IBIs (factor: time) were analyzed separately for hits and misses (factor: detection). The analysis showed significant main effects of time (Greenhouse–Geisser corrected; *F*(5, 160) = 57.9, *p* < 0.001, ε = 0.299, η^2^*_G_* = 0.008) and detection (*F*(1, 32) = 6.37, *p* = 0.020, η^2^*_G_* = 0.0004) as well as their significant interaction (Greenhouse–Geisser corrected *F*(5, 160) = 13.5, *p* < 0.001, ε = 0.399, η^2^*_G_* = 0.0003; Fig. 4A). IBIs prior to the stimulus (*S-1*, *S-2*) did not differ significantly between hits and misses. IBIs concurrent with the stimulus were significantly longer for hits than for misses (*Stimulus*: *t*(32) = 4.21, *p* = 0.006). This effect was also observed for IBIs right after the stimulus (*S+1*: *t*(32) = 5.22, *p* < 0.001) but not for subsequent IBIs (*S+2*, *S+3*). For both hits and misses, a significant cardiac deceleration was found between IBI before and during the stimulus (from *S-1* to *Stimulus*, hits: *t*(32) = −7.28, *p* < 0.001, misses: *t*(32) = −4.96, *p* < 0.001) as well as between the IBI during and after the stimulus (from *Stimulus* to *S+1*, hits: *t*(32) = −5.95, *p* < 0.001, misses: *t*(32) = −5.00, *p* < 0.001). Cardiac deceleration was followed by an immediate acceleration for hits (from *S+1* to *S+2*: *t*(32) = 4.93, *p* < 0.001) which was not observed for misses (from *S+1* to *S+2*: *t*(32) = 1.28, *p*_corrected_ = 1.000). In the later phase of the trials (from *S+2* to *S+3*) cardiac acceleration was present after both hits (*t*(32) = 7.50, *p* < 0.001) and misses (*t*(32) = 5.67, *p* < 0.001).

To further explore the association between cardiac deceleration and conscious somatosensory perception, the “slopes” of the stimulus-induced heartbeat deceleration (*Stimulus* - *S-1*) were compared between hits and misses. Conscious perception was accompanied by larger cardiac deceleration (*M* = 0.07, *SD* = 0.08) than missing the stimulus (*M* = −0.08, *SD* = 0.07), *t*(32) = 6.97, *p* < 0.001, Cohen’s *d* = 1.21 (Fig. 4B).

**Figure 4.**
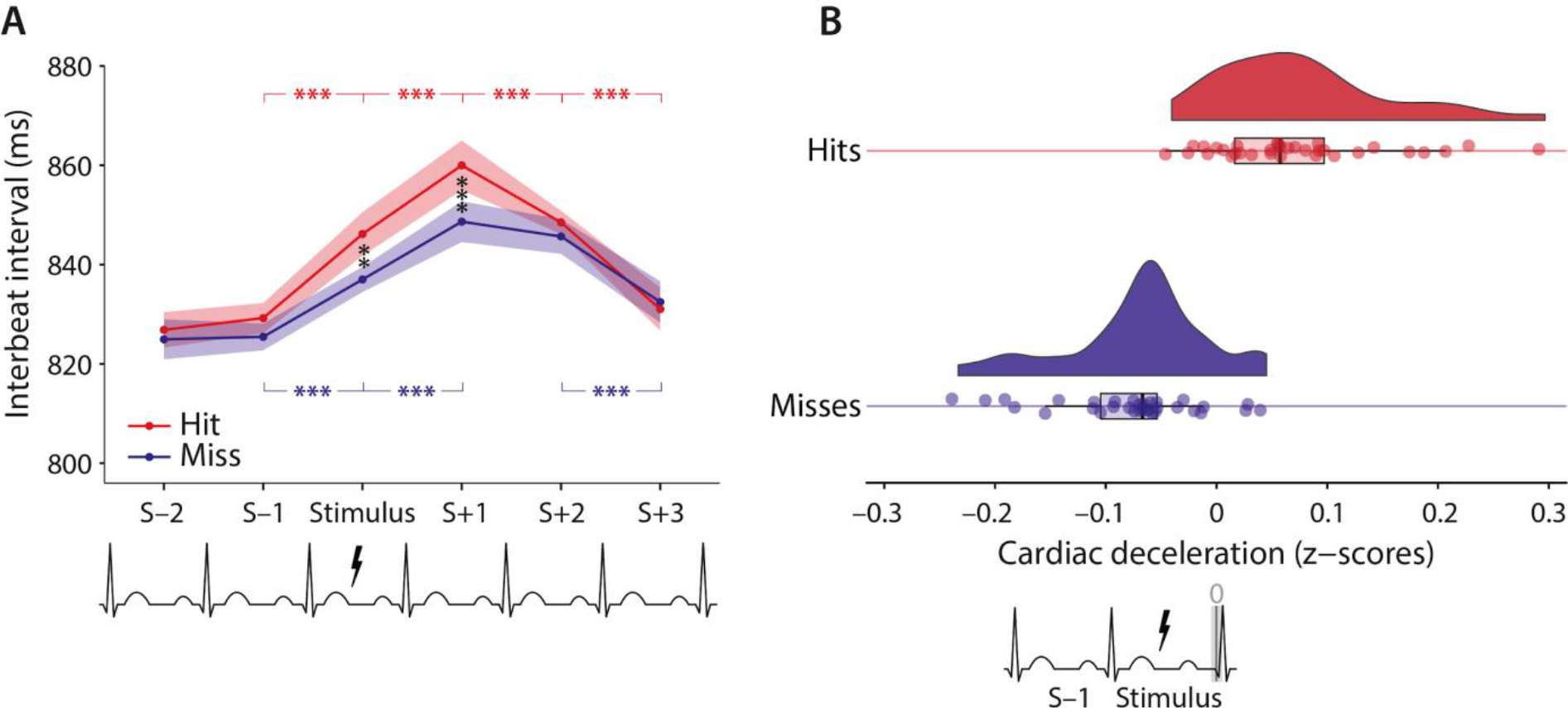
Association between heart rate and perceptual performance over the course of a trial. (**A**) Mean interbeat intervals (IBIs) for hits and misses: two IBIs before (*S-1*, *S-2*) and three IBIs after (*S+1*, *S+2*, *S+3*) the stimulus onset (*Stimulus*). In sum, the previously found deceleration-acceleration pattern was observed during the detection task, with more pronounced cardiac deceleration after hits than after misses. The vertical and horizontal bars with asterisks indicate significant pairwise post-hoc comparisons. (**B**) The extent of cardiac deceleration as a correlate of conscious perception, visualized using raincloud plots (Allen, Poggiali, Whitaker, Marshall, & Kievit, 2018). The standardized slope of cardiac deceleration (i.e., the difference between *Stimulus* and *S-1*) was greater for hits than for misses. Confidence bands indicate within-participants standard error of the mean, ** *p* < 0.01, *** *p* < 0.001.

## 4. DISCUSSION

In this study, we investigated if conscious somatosensory perception varies across the cardiac cycle and how it interacts with the heart rate. We found that the detection of near-threshold electrical finger nerve stimulation is significantly increased during diastole compared to systole. We also found (1) no evidence that the heart rate before a stimulus influenced perceptual performance, (2) conscious detection was associated with a stronger cardiac deceleration in the IBI during and after a stimulus, and (3) the heart rate accelerated with a delay after non-detected compared to detected stimuli. Taken together, these results indicate that conscious access to somatosensory signals varies across the cardiac cycle and transiently decreases the heart rate.

The difference between our findings and a previous study, which reported increased somatosensory sensitivity during “systole” (R+300 ms) compared to “diastole” (R+600 ms and R+0 ms; Edwards et al., 2009), may have several – also methodological – reasons: a) Edwards et al. (2009) considered R+0 ms as representative for diastole, while we – in the binary analysis – considered it as the beginning of systole; in addition, visual stimuli presented synchronously to the heartbeat (R peak) have been shown to be perceptually suppressed (Salomon et al., 2016) which hinders the interpretation of R-peak-related effects; b) in our analysis of detection across the entire cardiac cycle, near-threshold stimulus detection was increased in the very late phase of the cardiac cycle (beyond R+600 ms), which was not covered in the study by Edwards et al. (2009).

A possible physiological mechanism for the relatively increased detection during diastole is the baroreceptor-driven inhibition of sensory neural systems during systole (Duschek et al., 2013; Critchley & Garfinkel, 2015). This is consistent with previous findings, in which baroreceptor activity has been related to lower intensity ratings for acoustic (Cohen et al., 1980; Schulz et al., 2009) and painful stimuli (Wilkinson et al., 2013) as well as longer reaction times (Birren et al., 1963; Edwards et al., 2007; McIntyre et al., 2008) for stimuli presented early (i.e., at systole) compared to late (i.e., at diastole) in the cardiac cycle. However, there is also evidence that specifically threatening visual stimuli are perceived more easily and rated as more intense during systole (Garfinkel et al., 2014). Yet, the faint electrical stimulation in our study does not qualify as a threat signal but is rather an emotionally neutral stimulus, as they are typically used in studies of cardiac effects on conscious perception.

More broadly, it is not clear whether perceptual fluctuations related to rhythmic activity of the body and the brain (such as the heartbeat and respiration, and various forms of brain rhythms) come with an overall “functional advantage” or whether they are just an epiphenomenal consequence of physiological and anatomical constraints. For neural oscillations (e.g., alpha-band related variations in visual perception; Busch, Dubois, & VanRullen, 2009; Dugué, Marque, & VanRullen, 2011), it remains a matter of debate how perception benefits from inherent rhythmicity (VanRullen, 2016). It has been proposed that brain oscillations serve the effective communication between neurons (Fries, 2015) and enable the simultaneous encoding of multiple stimulus features (Lisman, 2005; VanRullen et al., 2005). However, perceptual rhythms in the brain have also been suggested to not have any functional role but result from satisfying biological constraints (VanRullen, 2016). A similar point could be made about the role of cardiac-related fluctuations in perception – especially because both are likely to be linked (Klimesch, 2018; Klimesch; 2013).

The present findings could also be understood as suppressing weak and non-salient somatosensory signals from reaching consciousness during baroreceptor firing. Given the enhanced processing of threat stimuli (Garfinkel et al., 2014) and pain inhibition during systole (Wilkinson et al. 2013), it has been proposed that baroreceptor signals promote a “fight-or-flight” mode of behavior (Garfinkel & Critchley, 2016). In line with this interpretation, Pramme et al. (2016) reported enhanced visual selection during systole. Hence, baroreceptor-mediated inhibition of cortical activation might facilitate the allocation of attention to situationally relevant stimuli (Pramme, Larra, Schächinger, & Frings, 2016). It could be hypothesized that a stressor-evoked heart rate increase facilitates the processing of situation-relevant information in the external world; by shortening diastole rather than systole (Herzog et al, 2002) this results in proportionally longer periods, during which non-salient stimuli are inhibited. Future studies could explore the functional role of perceptual periodicity, for example by manipulating the salience of the near-threshold signals through different task requirements or an association with threatening stimuli (e.g., declaring or animating near-threshold somatosensory stimuli as bites from Malaria-infected mosquitoes).

Further, accounting for the bidirectional information flow between the heart and the brain (Faes et al., 2017, Lin et al., 2016), we investigated the influence of the (pre-stimulus) heart rate on perception and, in turn, the influence of perception on (post-stimulus) heart rate changes. Even though it was early hypothesized that cardiac deceleration enhances perceptual sensitivity (Graham & Clifton, 1965; Lacey et al., 1963; Lacey, 1967; Sandman, 1986; but see also Elliot, 1972), results are inconsistent in the auditory (Edwards & Alsip, 1969, Saxon & Dahle, 1971) or visual (Cobos et al., 2018; McCanne & Sandman, 1974, Park et al., 2014, Sandman et al., 1977) and outright lacking in the somatosensory modality. Our findings match reports in the visual domain (Park et al., 2014; Cobos et al., 2018) with respect to (1) the lack of evidence for the influence of the pre-stimulus heart rate on detection and (2) a more pronounced cardiac deceleration after detecting (relative to not detecting) near-threshold stimuli. In addition, we found that also the interbeat interval length *during* the somatosensory stimulation differed between hits and misses, and that the extent of cardiac deceleration coincident with the stimulation was higher for detected compared to non-detected stimuli. Moreover, for non-detected stimuli, we observed a delayed cardiac acceleration, which might be a side effect of less pronounced deceleration after misses, but has been also reported to occur after an incorrect visual stimuli discrimination (Łukowska, Sznajder, & Wierzchoń, 2018) and, more broadly, is thought to reflect the processing of erroneous responses (Crone et al., 2003; Danev & de Winter, 1971; Fiehler, Ullsperger, Grigutsch, & Cramon, 2004; Hajcak, McDonald, & Simons, 2003).

This lengthening of the interbeat interval reflects the rapid autonomic (i.e., parasympathetic) response to the consciously perceived stimulus (Barry, 2006; Knippenberg, Barry, Kuniecki, & van Luijtelaar, 2012). Due to its speed, it is likely to also affect the duration of the interbeat interval, during which the stimulus is presented (Lacey & Lacey, 1977; Jennings, van der Molen, Somsen, & Brock, 1991; Jennings & van der Molen, 1993; Velden, Barry, & Wölk, 1987; Zimmermann, Velden, & Wölk, 1991; but see also Barry, 1993). It could also be that both cardiac deceleration and enhanced detection are the result of the central processes responsible for attentional preparation, which involve the activity of inhibitory brain circuits (Aron et al., 2007). Particularly subthalamic nuclei have been proposed to regulate the extent of (preparatory) cardiac deceleration (Jennings, Molen, & Tanase, 2009), which – as a marker of increased vigilance (Barry, 1988, 1996) – has also been shown to predict accuracy in tasks requiring skilled motor performance (Fahimi & Vaezmousavi, 2011; Tremayne & Barry, 2001). Even though our design minimized the influence of preparation attempts by randomizing stimulus onsets, it cannot be ruled out that the concomitant increases in cardiac deceleration and conscious detection were both caused by coincident peaks of attentional engagement (Fiebelkorn & Kastner, 2018).

The present study has several limitations: First, as baroreceptor or brain activity were not directly measured, we can only speculate about the baroreceptor influences on (central) sensory processing. Second, the lack of significant differences in cardiac effects on somatosensory perception between the groups with high and low interoceptive accuracy might be due to the limited sample sizes or the insufficient validity of Heartbeat Counting Task itself (Brener & Ring, 2016). Third, to apply signal detection theory measures (Green & Swets, 1966), future studies should allow to temporally locate false alarms within the cardiac cycle. In the current design, we used a non-cued stimulus onset within a 1,800-ms time window. This precluded to determine the position of false alarms within the cardiac phases. Visual or acoustic cues for stimulus onsets would suffice for this purpose but may themselves introduce crossmodal interactions (Dionne, Meehan, Legon, & Staines, 2010), for which the influence of the cardiac cycle remains unknown. Future research could also consider including a graded measure of stimulus awareness (Ramsøy & Overgaard, 2004; Sandberg, Timmermans, Overgaard, & Cleeremans, 2010) to parametrically assess the effects of (un)conscious sensory processing on cardiac activity.

In sum, we find that conscious perception of somatosensory stimuli varies across the cardiac cycle and is associated with increased cardiac deceleration. This highlights the importance of activity in the autonomic nervous system for perceptual awareness. Our findings emphasize the irreducible relevance of bodily states for sensory processing and suggest a more holistic picture of an organism’s cognition, for which contributions from the brain and from the rest of the body cannot be clearly separated.

## AUTHOR NOTES

### Acknowledgments

We thank Sylvia Stasch for her assistance in data acquisition as well as Sven Ohl, Till Nierhaus, Birol Taskin, Fivos Iliopoulos, and Stella Kunzendorf for their methodological advice and useful discussions.

## REFERENCES

Abboud, F. M., & Chapleau, M. W. (1988). Effects of pulse frequency on single-unit baroreceptor activity during sine-wave and natural pulses in dogs. The Journal of Physiology, 401, 295–308. https://doi.org/10.1113/jphysiol.1988.sp017163

Agostinelli, C., & Lund, U. (2013) R package ‘circular’: Circular Statistics (version 0.4-7). Retrieved from URL: https://r-forge.r-project.org/projects/circular/

Allen, M., Poggiali, D., Whitaker, K., Marshall, T. R., & Kievit, R. (2018). Raincloud plots: a multi-platform tool for robust data visualization. Peer J Preprints 6:e27137v1. https://doi.org/10.7287/peerj.preprints.27137v1

Aron, A. R., Durston, S., Eagle, D. M., Logan, G. D., Stinear, C. M., & Stuphorn, V. (2007). Converging Evidence for a Fronto-Basal-Ganglia Network for Inhibitory Control of Action and Cognition. Journal of Neuroscience, 27(44), 11860–11864. https://doi.org/10.1523/JNEUROSCI.3644-07.2007

Barrett, C. J., & Bolter, C. P. (2006). The influence of heart rate on baroreceptor fibre activity in the carotid sinus and aortic depressor nerves of the rabbit. Experimental Physiology, 91(5), 845–852. https://doi.org/10.1113/expphysiol.2006.033902

Barrett, L. F., & Simmons, W. K. (2015). Interoceptive predictions in the brain. Nature Reviews. Neuroscience, 16(7), 419–429. https://doi.org/10.1038/nrn3950

Barry, R. J. (1988). Significance and components of the orienting response: effects of signal value versus vigilance. International Journal of Psychophysiology, 6(4), 343–346. https://doi.org/10.1016/0167-8760(88)90023-2

Barry, R. J. (1993). Graphical and statistical techniques for cardiac cycle time (phase) dependent changes in interbeat interval: Problems with the Jennings et al. (1991) proposals. Biological Psychology, 35(1), 59–65. https://doi.org/10.1016/0301-0511(93)90092-M

Barry, R. J. (1996). Preliminary process theory: towards an integrated account of the psychophysiology of cognitive processes. Acta Neurobiologiae Experimentalis, 56(1), 469–484.

Barry, R. J. (2006). Promise versus reality in relation to the unitary orienting reflex: A case study examining the role of theory in psychophysiology. International Journal of Psychophysiology, 62(3), 353–366. https://doi.org/10.1016/j.ijpsycho.2006.01.004

Birren, J. E., Cardon, P. V., & Phillips, S. L. (1963). Reaction time as a function of the cardiac cycle in young adults. Science, 140(3563), 195–196. https://dx.doi.org/10.1126/science.140.3563.195-a

Boudoulas, H., Geleris, P., Lewis, R. P., & Rittgers, S. E. (1981). Linear Relationship Between Electrical Systole, Mechanical Systole, and Heart Rate. Chest, 80(5), 613–617. https://doi.org/10.1378/chest.80.5.613

Börger, N., & Meere, J. van der. (2000). Motor control and state regulation in children with ADHD: A cardiac response study. Biological Psychology, 51(2–3), 247–267. https://doi.org/10.1016/S0301-0511(99)00040-X

Cobos, M. I., Guerra, P. M., Vila, J., & Chica, A. B. (2018). Heart-rate modulations reveal attention and consciousness interactions. Psychophysiology, e13295. https://doi.org/10.1111/psyp.13295

Bradley, M. M., Cuthbert, B. N., & Lang, P. J. (1990). Startle reflex modification: emotion or attention? Psychophysiology, 27(5), 513–522. https://doi:10.1111/j.1469-8986.1990.tb01966.x

Brainard, D. H. (1997). The Psychophysics Toolbox. Spatial Vision, 10(4), 433–436. https://doi.org/10.1163/156856897X00357

Brener, J., & Ring, C. (2016). Towards a psychophysics of interoceptive processes: the measurement of heartbeat detection. Philosophical Transactions of the Royal Society of London. Series B, Biological Sciences, 371(1708). https://doi.org/10.1098/rstb.2016.0015

Busch, N. A., Dubois, J., & VanRullen, R. (2009). The phase of ongoing EEG oscillations predicts visual perception. The Journal of Neuroscience: The Official Journal of the Society for Neuroscience, 29(24), 7869–7876. https://doi.org/10.1523/JNEUROSCI.0113-09.2009

Coblentz, B., Harvey, R. M., Ferrer, M. I., Cournand, A., & Richards, D. W. (1949). The Relationship Between Electrical and Mechanical Events in the Cardiac Cycle of Man. Heart, 11(1), 1–22. https://doi.org/10.1136/hrt.11.1.1

Cohen, R., Lieb, H., & Rist, F. (1980). Loudness judgments, evoked potentials, and reaction time to acoustic stimuli early and late in the cardiac cycle in chronic schizophrenics. Psychiatry Research, 3(1), 23–29. https://doi.org/10.1016/0165-1781(80)90044-X

Christofaro, D. G. D., Casonatto, J., Vanderlei, L. C. M., Cucato, G. G., & Dias, R. M. R. (2017). Relationship between Resting Heart Rate, Blood Pressure and Pulse Pressure in Adolescents. Arquivos Brasileiros de Cardiologia, 108(5), 405–410. https://doi.org/10.5935/abc.20170050

Critchley, H. D., & Garfinkel, S. N. (2015). Interactions between visceral afferent signaling and stimulus processing. Frontiers in Neuroscience, 9. https://doi.org/10.3389/fnins.2015.00286

Critchley, H. D., & Garfinkel, S. N. (2017). Interoception and emotion. Current Opinion in Psychology, 17, 7–14. https://doi.org/10.1016/j.copsyc.2017.04.020

Critchley, H. D., & Garfinkel, S. N. (2018). The influence of physiological signals on cognition. Current Opinion in Behavioral Sciences, 19, 13–18. https://doi.org/10.1016/j.cobeha.2017.08.014

Craig, A. D. B. (2009). How do you feel – now? The anterior insula and human awareness. Nature Reviews. Neuroscience, 10(1), 59–70. https://doi.org/10.1038/nrn2555

Crone, E. A., Van der Veen, F. M., Van der Molen, M.W., Somsen, R. J., Van Beek, B., & Jennings, J. (2003). Cardiac concomitants of feedback processing. Biological Psychology, 64(1–2), 143–156. https://doi.org/10.1016/S0301-0511(03)00106-6

Danev, S. G., & de Winter, C. R. (1971). Heart rate deceleration after erroneous responses. Psychologische Forschung, 35(1), 27–34. https://doi.org/10.1007/BF00424472

Dampney, R. A. L. (2016). Central neural control of the cardiovascular system: current perspectives. Advances in Physiology Education, 40(3), 283–296. https://doi.org/10.1152/advan.00027.2016

Davis, R. C., & Buchwald, A. M. (1957). An exploration of somatic Response patterns: Stimulus and sex differences. Journal of Comparative and Physiological Psychology, 50(1), 44–52. https://doi.org/10.1037/h0046428

Davis, R. C., Buchwald, A. M., & Frankmann, R. W. (1955). Autonomic and muscular responses, and their relation to simple stimuli. Psychological Monographs: General and Applied, 69(20), 1–71. https://doi.org/10.1037/h0093734

Delfini, L. F., & Campos, J. J. (1972). Signal Detection and the “Cardiac Arousal Cycle.” Psychophysiology, 9(5), 484–491. https://doi.org/10.1111/j.1469-8986.1972.tb01801.x

Di Lernia, D., Serino, S., Pezzulo, G., Pedroli, E., Cipresso, P., & Riva, G. (2018). Feel the Time. Time Perception as a Function of Interoceptive Processing. Frontiers in Human Neuroscience, 12, 74. https://doi.org/10.3389/fnhum.2018.00074

Dionne, J. K., Meehan, S. K., Legon, W., & Staines, W. R. (2010). Crossmodal influences in somatosensory cortex: Interaction of vision and touch. Human Brain Mapping, 31(1), 14–25. https://doi.org/10.1002/hbm.20841

Dugué, L., Marque, P., & VanRullen, R. (2011). The Phase of Ongoing Oscillations Mediates the Causal Relation between Brain Excitation and Visual Perception. Journal of Neuroscience, 31(33), 11889–11893. https://doi.org/10.1523/JNEUROSCI.1161-11.2011

Dunn, B. D., Galton, H. C., Morgan, R., Evans, D., Oliver, C., Meyer, M., … Dalgleish, T. (2010). Listening to Your Heart: How Interoception Shapes Emotion Experience and Intuitive Decision Making. Psychological Science, 21(12), 1835–1844. https://doi.org/10.1177/0956797610389191

Duschek, S., Werner, N. S., & Reyes Del Paso, G. A. (2013). The behavioral impact of baroreflex function: a review. Psychophysiology, 50(12), 1183–1193. https://doi.org/10.1111/psyp.12136

Edwards, D. C., & Alsip, J. E. (1969). Stimulus Detection During Periods of High and Low Heart Rate. Psychophysiology, 5(4), 431–434. https://doi.org/10.1111/j.1469-8986.1969.tb02843.x

Edwards, L., Ring, C., McIntyre, D., Carroll, D., & Martin, U. (2007). Psychomotor speed in hypertension: Effects of reaction time components, stimulus modality, and phase of the cardiac cycle. Psychophysiology, 44(3), 459–468. https://doi.org/10.1111/j.1469-8986.2007.00521.x

Edwards, L., Ring, C., McIntyre, D., Winer, J. B., & Martin, U. (2009). Sensory detection thresholds are modulated across the cardiac cycle: evidence that cutaneous sensibility is greatest for systolic stimulation. Psychophysiology, 46(2), 252–256. https://doi.org/10.1111/j.1469-8986.2008.00769.x

Elliott, R. (1972). The significance of heart rate for behavior: A critique of Lacey’s hypothesis. Journal of Personality and Social Psychology, 22(3), 398–409. https://doi.org/10.1037/h0032832

Elliott, R., & Graf, V. (1972). Visual Sensitivity as a Function of Phase of Cardiac Cycle. Psychophysiology, 9(3), 357–361. https://doi.org/10.1111/j.1469-8986.1972.tb03219.x

Faes, L., Greco, A., Lanata, A., Barbieri, R., Scilingo, E. P., & Valenza, G. (2017). Causal brain-heart information transfer during visual emotional elicitation in healthy subjects: Preliminary evaluations and future perspectives. Conference Proceedings: … Annual International Conference of the IEEE Engineering in Medicine and Biology Society. IEEE Engineering in Medicine and Biology Society. Annual Conference, 2017, 1559–1562. https://doi.org/10.1109/EMBC.2017.8037134

Fahimi, F., & Vaezmousavi, M. (2011). Physiological Patterning of Short Badminton Serve: A Psychophysiological Perspective to Vigilance and Arousal. World Applied Sciences Journal, 12, 347–353.

Fiebelkorn, I. C., & Kastner, S. (2018). A Rhythmic Theory of Attention. Trends in Cognitive Sciences. https://doi.org/10.1016/j.tics.2018.11.009

Fiehler, K., Ullsperger, M., Grigutsch, M., & von Cramon, D. Y. V. (2004). Cardiac responses to error processing and response conflict. In M. Ullsperger & M. Falkenstein (Eds.), Errors, conflicts, and the brain. Current opinions on performance monitoring (pp. 135–140). Leipzig: Max Planck Institute for Human Cognitive and Brain Sciences.

Fridericia, L. S. (1921). Die Systolendauer im Elektrokardiogramm bei normalen Menschen und bei Herzkranken. Acta Medica Scandinavica, 54(1), 17–50. https://doi.org/10.1111/j.0954-6820.1921.tb15167.x

Fries, P. (2015). Rhythms for Cognition: Communication through Coherence. Neuron, 88(1), 220–235. https://doi.org/10.1016/j.neuron.2015.09.034

Fukushima, H., Terasawa, Y., & Umeda, S. (2011). Association between interoception and empathy: evidence from heartbeat-evoked brain potential. International Journal of Psychophysiology, 79(2), 259–265. https://doi.org/10.1016/j.ijpsycho.2010.10.015

Garfinkel, S. N., & Critchley, H. D. (2016). Threat and the Body: How the Heart Supports Fear Processing. Trends in Cognitive Sciences, 20(1), 34–46. https://doi.org/10.1016/j.tics.2015.10.005

Garfinkel, S. N., Seth, A. K., Barrett, A. B., Suzuki, K., & Critchley, H. D. (2015). Knowing your own heart: distinguishing interoceptive accuracy from interoceptive awareness. Biological Psychology, 104, 65–74. https://doi.org/10.1016/j.biopsycho.2014.11.004

Garfinkel, S. N., Minati, L., Gray, M. A., Seth, A. K., Dolan, R. J., & Critchley, H. D. (2014). Fear from the Heart: Sensitivity to Fear Stimuli Depends on Individual Heartbeats. The Journal of Neuroscience, 34(19), 6573–6582. https://doi.org/10.1523/JNEUROSCI.3507-13.2014

Gill, H., & Hoffmann, A. K. (2010). The timing of onset of mechanical systole and diastole in reference to the QRS-T complex: a study to determine performance criteria for a non-invasive diastolic timed vibration massage system in treatment of potentially unstable cardiac disorders. Cardiovascular Engineering, 10(4), 235–245. https://doi.org/10.1007/s10558-010-9108-x

Green, D. M., & Swets, J. A. (1966). Signal detection theory and psychophysics. Oxford, England: John Wiley.

Grynberg, D., & Pollatos, O. (2015). Perceiving one’s body shapes empathy. Physiology & Behavior, 140, 54–60. https://doi.org/10.1016/j.physbeh.2014.12.026

Gu, X., & FitzGerald, T. H. B. (2014). Interoceptive inference: homeostasis and decision-making. Trends in Cognitive Sciences, 18(6), 269–270. https://doi.org/10.1016/j.tics.2014.02.001

Graham, F. K., & Clifton, R. K. (1966). Heart-rate change as a component of the orienting response. Psychological Bulletin, 65(5), 305–320. https://doi.org/10.1037/h0023258

Gray, M. L., & Crowell, D. H. (1968). Heart rate changes to sudden peripheral stimuli in the human during early infancy. The Journal of Pediatrics, 72(6), 807–814. https://doi.org/10.1016/S0022-3476(68)80433-0

Greenwald, M. K., Cook, E. W., & Lang, P. J. (1989). Affective judgment and psychophysiological response: Dimensional covariation in the evaluation of pictorial stimuli. Journal of Psychophysiology, 3(1), 51–64.

Hajcak, G., McDonald, N., & Simons, R. F. (2003). To err is autonomic: error-related brain potentials, ANS activity, and post-error compensatory behavior. Psychophysiology, 40(6), 895–903. https://doi.org/10.1111/1469-8986.00107

Hare, R. D. (1973). Orienting and Defensive Responses to Visual Stimuli. Psychophysiology, 10(5), 453–464. https://doi.org/10.1111/j.1469-8986.1973.tb00532.x

Herzog, C., Abolmaali, N., Balzer, J. O., Baunach, S., Ackermann, H., Doğan, S., … Vogl, T. J. (2002). Heart-rate-adapted image reconstruction in multidetector-row cardiac CT: influence of physiological and technical prerequisite on image quality. European Radiology, 12, 2670–2678. https://doi.org/10.1007/s00330-002-1553-5

Jennings, J. R., & van der Molen, M. W. (1993). Phase sensitivity: A dead end or a challenge? Biological Psychology, 35, 67–71. https://doi.org/10.1016/0301-0511(93)90093-n

Jennings, J. R., van der Molen, M. W., & Tanase, C. (2009). Preparing hearts and minds: Cardiac slowing and a cortical inhibitory network. Psychophysiology, 46(6), 1170–1178. https://doi.org/10.1111/j.1469-8986.2009.00866.x

Jennings, J. R., van der Molen, M. W., Somsen, R. J., & Brock, K. (1991). Weak sensory stimuli induce a phase sensitive bradycardia. Psychophysiology, 28(1), 1–10. https://doi.org/10.1111/j.1469-8986.1991.tb03380.x

Kingdom, F. A. A., & Prins, N. (2010). Psychophysics: A practical introduction. San Diego, CA, US: Elsevier Academic Press.

Kleckner, I. R., Zhang, J., Touroutoglou, A., Chanes, L., Xia, C., Simmons, W. K., … Barrett, L. F. (2017). Evidence for a Large-Scale Brain System Supporting Allostasis and Interoception in Humans. Nature Human Behaviour, 1. https://doi.org/10.1038/s41562-017-0069

Kleiner, M., Brainard, D., Pelli, D., Ingling, A., Murray, R., & Broussard, C. (2007). What’s new in psychtoolbox-3. Perception, 36(14), 1–16. https://doi.org/10.1068/v070821

Klimesch, W. (2013). An algorithm for the EEG frequency architecture of consciousness and brain body coupling. Frontiers in Human Neuroscience, 7. https://doi.org/10.3389/fnhum.2013.00766

Klimesch, W. (2018). The frequency architecture of brain and brain body oscillations: an analysis. European Journal of Neuroscience, 48(7), 2431–2453. https://doi.org/10.1111/ejn.14192

Knippenberg, J. M. J., Barry, R. J., Kuniecki, M. J., & van Luijtelaar, G. (2012). Fast, transient cardiac accelerations and decelerations during fear conditioning in rats. Physiology & Behavior, 105(3), 607–612. https://doi.org/10.1016/j.physbeh.2011.09.018

Lacey, J. (1967). Somatic response patterning and stress: some revisions of activation theory. In M. II. Appley & R. Trumball (Eds.), Psychological stress: Issues in Research (pp. 14–42). New York: Appleton-Century-Crofts.

Lacey, J. I., & Lacey, B. C. (1970) Some autonomic-central nervous system interrelationships. In P. Black (Ed.), Physiological correlates of emotion (pp. 205–227). New York: Academic Press.

Lacey, J. I., & Lacey, B. C. (1974) On heart rate responses and behavior: A reply to Elliott. Journal of Personality and Social Psychology, 30(1), 1–18. http://dx.doi.org/10.1037/h0036559

Lacey, B. C., & Lacey, J. I. (1977). Change in heart period: A function of sensorimotor event timing within the cardiac cycle. Physiological Psychology, 5(3), 383–393. https://doi.org/10.3758/BF03335349

Lacey, J. I., Kagan, J., Lacey, B. C., & Moss, H. A. (1963). The visceral level: Situational determinants and behavioral correlates of autonomic response patterns. In P. H. Knapp (Ed.), Expressions of the emotions in man (pp. 161–196). New York: International Universities Press.

Landgren, W. (1952). On the Excitation Mechanism of the Carotid Baroceptors. Acta Physiologica Scandinavica, 26(1), 1–34. https://doi.org/10.1111/j.1748-1716.1952.tb00889.x

Lewis, R. L., Rittogers, S. E., Froester, W. F., & Boudoulas, H. (1977). A critical review of the systolic time intervals. Circulation, 56(2), 146–158.

Libby, W. L., Lacey, B. C., & Lacey, J. I. (1973). Pupillary and Cardiac Activity During Visual Attention. Psychophysiology, 10(3), 270–294. https://doi.org/10.1111/j.1469-8986.1973.tb00526.x

Lin, A., Liu, K., Bartsch, R., & Ivanov, P. (2016). Delay-correlation landscape reveals characteristic time delays of brain rhythms and heart interactions. Philosophical Transactions of the Royal Society A: Mathematical, Physical and Engineering Sciences, 374(2067). https://doi.org/10.1098/rsta.2015.0182

Lisman, J. (2005). The theta/gamma discrete phase code occuring during the hippocampal phase precession may be a more general brain coding scheme. Hippocampus, 15(7), 913–922. https://doi.org/10.1002/hipo.20121

Łukowska, M., Sznajder, M., & Wierzchoń, M. (2018). Error-related cardiac response as information for visibility judgements. Scientific Reports, 8(1), 1131. https://doi.org/10.1038/s41598-018-19144-0

Makovac, E., Garfinkel, S. N., Bassi, A., Basile, B., Macaluso, E., Cercignani, M., … Critchley, H. (2015). Effect of Parasympathetic Stimulation on Brain Activity During Appraisal of Fearful Expressions. Neuropsychopharmacology, 40(7), 1649–1658. https://doi.org/10.1038/npp.2015.10

Mancia, G., & Mark, A. L. (2011). Arterial Baroreflexes in Humans. In R. Terjung (Ed.), Comprehensive Physiology. Hoboken, NJ, USA: John Wiley & Sons, Inc. https://doi.org/10.1002/cphy.cp020320

Mancia, G., Ferrari, A., Gregorini, L., Parati, G., Pomidossi, G., Bertinieri, G., … Zanchetti, A. (1983). Blood pressure and heart rate variabilities in normotensive and hypertensive human beings. Circulation Research, 53(1), 96–104. https://doi.org/10.1161/01.RES.53.1.96

McCanne, T. R., & Sandman, C. A. (1974). Instrumental Heart Rate Responses and Visual Perception: A Preliminary Study. Psychophysiology, 11(3), 283–287. https://doi.org/10.1111/j.1469-8986.1974.tb00545.x

McIntyre, D., Ring, C., Edwards, L., & Carroll, D. (2008). Simple reaction time as a function of the phase of the cardiac cycle in young adults at risk for hypertension. Psychophysiology, 45(2), 333–336. https://doi.org/10.1111/j.1469-8986.2007.00619.x

Meissner, K., & Wittmann, M. (2011). Body signals, cardiac awareness, and the perception of time. Biological Psychology, 86(3), 289–297. https://doi.org/10.1016/j.biopsycho.2011.01.001

Otten, L. J., Gaillard, A. W. K., & Wientjes, C. J. E. (1995). The relation between event-related brain potential, heart rate, and blood pressure responses in an S1-S2 paradigm. Biological Psychology, 39(2), 81–102. https://doi.org/10.1016/0301-0511(94)00969-5

Park, H.-D., & Tallon-Baudry, C. (2014). The neural subjective frame: from bodily signals to perceptual consciousness. Philosophical Transactions of the Royal Society of London. Series B, Biological Sciences, 369(1641), 20130208. https://doi.org/10.1098/rstb.2013.0208

Park, H.-D., Correia, S., Ducorps, A., & Tallon-Baudry, C. (2014). Spontaneous fluctuations in neural responses to heartbeats predict visual detection. Nature Neuroscience, 17(4), 612–618. https://doi.org/10.1038/nn.3671

Pelli, D. G. (1997). The VideoToolbox software for visual psychophysics: transforming numbers into movies. Spatial Vision, 10(4), 437–442. https://doi.org/10.1163/156856897X00366

Pewsey, A., Neuhäuser, M., & Ruxton, G.D. (2013). Circular statistics in R. Oxford and New York: Oxford University Press.

Pramme, L., Larra, M. F., Schächinger, H., & Frings, C. (2016). Cardiac cycle time effects on selection efficiency in vision. Psychophysiology, 53(11), 1702–1711. https://doi.org/10.1111/psyp.12728

R Core Team (2016) R: A Language and Environment for Statistical Computing. R Foundation for Statistical Computing, Vienna. Retrieved from URL: https://www.r-project.org

Ramsøy, T. Z., & Overgaard, M. (2004). Introspection and subliminal perception. Phenomenology and the Cognitive Sciences, 3(1), 1–23

RStudio Team (2016). RStudio: Integrated Development Environment for R. Boston, MA: RStudio, Inc. Retrieved from URL: https://www.rstudio.com

Rau, H., & Elbert, T. (2001). Psychophysiology of arterial baroreceptors and the etiology of hypertension. Biological Psychology, 57(1–3), 179–201.

Rau, H., Pauli, P., Brody, S., Elbert, T., & Birbaumer, N. (1993). Baroreceptor stimulation alters cortical activity. Psychophysiology, 30(3), 322–325.

Réquin, J., & Brouchon, M. (1964). Mise en évidence chez l’ homme d’une fluctuation des seuils perceptifs visuels dans la période cardiaque. Comptes Rendus Des Séances de La Société de Biologie et de Ses Filiales, 158, 1891–1894.

Salomon, R., Ronchi, R., Dönz, J., Bello-Ruiz, J., Herbelin, B., Martet, R., … Blanke, O. (2016). The Insula Mediates Access to Awareness of Visual Stimuli Presented Synchronously to the Heartbeat. Journal of Neuroscience, 36(18), 5115–5127. https://doi.org/10.1523/JNEUROSCI.4262-15.2016

Sandberg, K., Timmermans, B., Overgaard, M., & Cleeremans, A. (2010). Measuring consciousness: Is one measure better than the other? Consciousness and Cognition, 19(4), 1069–1078. https://doi.org/10.1016/j.concog.2009.12.013

Sandman, C. A. (1986). Cardiac Afferent Influences on Consciousness. In R. J. Davidson, G. E. Schwartz, & D. Shapiro (Eds.), Consciousness and Self-Regulation: Advances in Research and Theory Volume 4 (pp. 55–85). Boston, MA: Springer US. https://doi.org/10.1007/978-1-4757-0629-1_3

Sandman, C. A., McCanne, T. R., Kaiser, D. N., & Diamond, B. (1977). Heart rate and cardiac phase influences on visual perception. Journal of Comparative and Physiological Psychology, 91(1), 189–202. http://dx.doi.org/10.1037/h0077302

Saxon, S. A. (1970). Detection of near threshold signals during four phases of cardiac cycle. The Alabama Journal of Medical Sciences, 7(4), 427–430.

Saxon, S. A., & Dahle, A. J. (1971). Auditory threshold variations during periods of induced high and low heart rates. Psychophysiology, 8(1), 23–29. https://doi.org/10.1111/j.1469-8986.1971.tb00433.x

Schulz, A., Reichert, C. F., Richter, S., Lass-Hennemann, J., Blumenthal, T. D., & Schächinger, H. (2009). Cardiac modulation of startle: effects on eye blink and higher cognitive processing. Brain and Cognition, 71(3), 265–271. https://doi.org/10.1016/j.bandc.2009.08.00

Seth, A. K. (2014). Response to Gu and FitzGerald: Interoceptive inference: from decision-making to organism integrity. Trends in Cognitive Sciences, 18(6), 270–271. https://doi.org/10.1016/j.tics.2014.03.006

Schandry, R. (1981) Heart Beat Perception and Emotional Experience. Psychophysiology, 18, 483–488. https://doi.org/10.1111/j.1469-8986.1981.tb02486.

Simons, R.F. (1988). Event-related slow brain potentials: A perspective from ANS psychophysiology. In: P.K. Ackles, J. R. Jennings, & M.G.H. Coles (Eds.), Advances in Psychophysiology, Vol. 3 (pp. 223–267). Greenwich: JAI Press Inc.

Uno, T., & Grings, W. W. (1965). Autonomic Components of Orienting Behavior. Psychophysiology, 1(4), 311–321. https://doi.org/10.1111/j.1469-8986.1965.tb03263.x

Suzuki, K., Garfinkel, S. N., Critchley, H. D., & Seth, A. K. (2013). Multisensory integration across exteroceptive and interoceptive domains modulates self-experience in the rubber-hand illusion. Neuropsychologia, 51(13), 2909–2917. https://doi.org/10.1016/j.neuropsychologia.2013.08.014

Tarvainen, M. P., Niskanen, J.‐P., Lipponen, J. A., Ranta‐aho, P. O., & Karjalainen, P. A. (2014). Kubios HRV – Heart rate variability analysis software. Computer Methods and Programs in Biomedicine, 113(1), 210–220. https://doi.org/10.1016/j.cmpb.2013.07.024

Tremayne, P., & Barry, R. J. (2001). Elite pistol shooters: physiological patterning of best vs. worst shots. International Journal of Psychophysiology, 41(1), 19–29. https://doi.org/10.1016/S0167-8760(00)00175-6

Wallace, A. G., Mitchell, J. H., Skinner, S. N., & Sarnoff, S. J. (1963). Duration of the Phases of Left Ventricular Systole. Circulation Research, 12(6), 611–619. https://doi.org/10.1161/01.RES.12.6.611

Walker, B. B., & Sandman, C. A. (1977). Physiological response patterns in ulcer patients: Phasic and tonic components of the electrogastrogram. Psychophysiology, 14(4), 393–400. https://doi.org/10.1111/j.1469-8986.1977.tb02971.x

Wiens, S. (2005). Interoception in emotional experience. Current Opinion in Neurology, 18(4), 442–7. https://doi.org/10.1097/01.wco.0000168079.92106.99

Wilkinson, M., McIntyre, D., & Edwards, L. (2013). Electrocutaneous pain thresholds are higher during systole than diastole. Biological Psychology, 94(1), 71–73. https://doi.org/10.1016/j.biopsycho.2013.05.002

Wilson, R. S. (1964). Autonomic changes produced by noxious and innocuous stimulation. Journal of Comparative and Physiological Psychology, 58(2), 290–295. https://doi:10.1037/h0045264

Wölk, C., Velden, M., Zimmermann, U., & Krug, S. (1989). The interrelation between phasic blood pressure and heart rate changes in the context of the “baroreceptor hypothesis.” Journal of Psychophysiology, 3(4), 397–402.

Valenza, G., Toschi, N., & Barbieri, R. (2016). Uncovering brain–heart information through advanced signal and image processing. Philosophical Transactions. Series A, Mathematical, Physical, and Engineering Sciences, 374(2067). https://doi.org/10.1098/rsta.2016.0020

VanRullen, R., Guyonneau, R., & Thorpe, S. J. (2005). Spike times make sense. Trends in Neurosciences, 28(1), 1–4. https://doi.org/10.1016/j.tins.2004.10.010

VanRullen, R. (2016). Perceptual Cycles. Trends in Cognitive Sciences, 20(10), 723–735. https://doi.org/10.1016/j.tics.2016.07.006

Vázquez-Seisdedos, C. R., Neto, J. E., Marañón Reyes, E. J., Klautau, A., & Limão de Oliveira, R. C. (2011). New approach for T-wave end detection on electrocardiogram: Performance in noisy conditions. BioMedical Engineering OnLine, 10(1), 77. https://doi.org/10.1186/1475-925X-10-77

Velden, M., & Juris, M. (1975). Perceptual Performance as a Function of Intra-Cycle Cardiac Activity. Psychophysiology, 12(6), 685–692. https://doi.org/10.1111/j.1469-8986.1975.tb00075.x

Velden, M., Barry, R. J., & Wölk, C. (1987). Time-dependent bradycardia: a new effect? International Journal of Psychophysiology, 4(4), 299–306. https://doi.org/10.1016/0167-8760(87)90042-0

Zimmerman, U., Velden, M., & Wölk, C. (1991). Empirical evidence against the “cycle time dependency” assumption. International Journal of Psychophysiology, 11(2), 125–129. https://doi.org/10.1016/0167-8760(91)90004-H

