## Supplementary Material for "Interactions between cardiac activity and conscious somatosensory perception"

### 1. SUPPLEMENTARY METHODS

In an exploratory analysis, we tested whether the link between conscious somatosensory perception and cardiac activity is influenced by the capacity to consciously perceive one's heartbeat (i.e., interoceptive accuracy; Garfinkel et al., 2015). After the completion of three blocks of somatosensory perception (and after the steel wire ring electrodes had been removed from the left index finger), interoceptive accuracy (Garfinkel et al., 2015) was measured using a Heartbeat Counting Task (Schandry, 1981): The participants were asked to count their heartbeats silently during acoustically signaled time intervals of the following lengths (presented in this order): 25, 45, 15, 55, and 35 s. Individual interoceptive accuracy (IA) scores were calculated comparing the objectively measured number of heartbeats and the subjectively reported number of heartbeats from the five intervals with the following formula: $1/5\ \Sigma\ (1 - (|recorded\ heartbeats - reported\ heartbeats|)/recorded\ heartbeats)$. The score varies between 0 and 1, with 1 indicating absolute accuracy of heartbeat perception. One participant was excluded due to not properly reported heartbeats (all zeroes). Also, one participant was excluded from the circular analyses due to non-uniformly distributed stimulation onsets across the cardiac cycle (see section 2.5 in the main text of the paper). In total, the supplementary circular analysis (section 2.2) included data from 31 participants (15 females, mean age = 25.7, *SD* = 4.0, range: 19-36 years) whereas the rest of supplementary analyses (sections: 2.1, 2.3, 2.4) included data from 32 participants (16 females, mean age = 25.8, *SD* = 4.0, range: 19-36 years).

### 2. SUPPLEMENTARY RESULTS

#### 2.1 Summary statistics for interoceptive accuracy

The mean IA score of 32 participants was 0.72 (*SD* = 0.15, range = 0.38-0.96). For further analyses, participants were median-split (*Me* = 0.75) into a high and a low IA group.

#### 2.2. Distribution of hits and misses across the cardiac cycle for high and low interoceptive accuracy groups

A Watson's Two Sample test was used to assess whether the angular directions of hit and miss concentration across the cardiac cycle differed between the groups of high and low IA, (Pewsey et al., 2013, p. 144). No significant group differences were found both for hits ($U^2$ = 0.09, $p$ > 0.05) or misses ($U^2$ = 0.07, $p$ > 0.05). Hence, there is no evidence that both groups differ with respect to the position of increased perceptual awareness for near-threshold somatosensory stimuli within the cardiac cycle. Further, Rayleigh tests were used to test the uniformity of hits and misses distribution across the cardiac cycle separately for high and low IA groups. There was no evidence for the non-uniformity of hits distribution both for high IA ($\overline{R}$ = 0.34, $p$ = 0.161) and low IA group ($\overline{R}$ = 0.38, $p$ = 0.114; Fig. S1A). Similarly for misses, the tests did not yield significant results both for high IA ($\overline{R}$ = 0.29, $p$ = 0.257) and low IA group ($\overline{R}$ = 0.42, $p$ = 0.072; Fig. S1B). These results suggest that interoceptive accuracy does not modulate the effects of the cardiac cycle on conscious somatosensory perception. Yet, the small sample sizes in the IA groups limit the statistical power of the used tests. As a side remark, continuous IA were not significantly correlated with the levels of hit ($r$(29) = 0.04, $p$ = 0.801) or miss concentration ($r$(29)= -0.13, $p$ = 0.485).

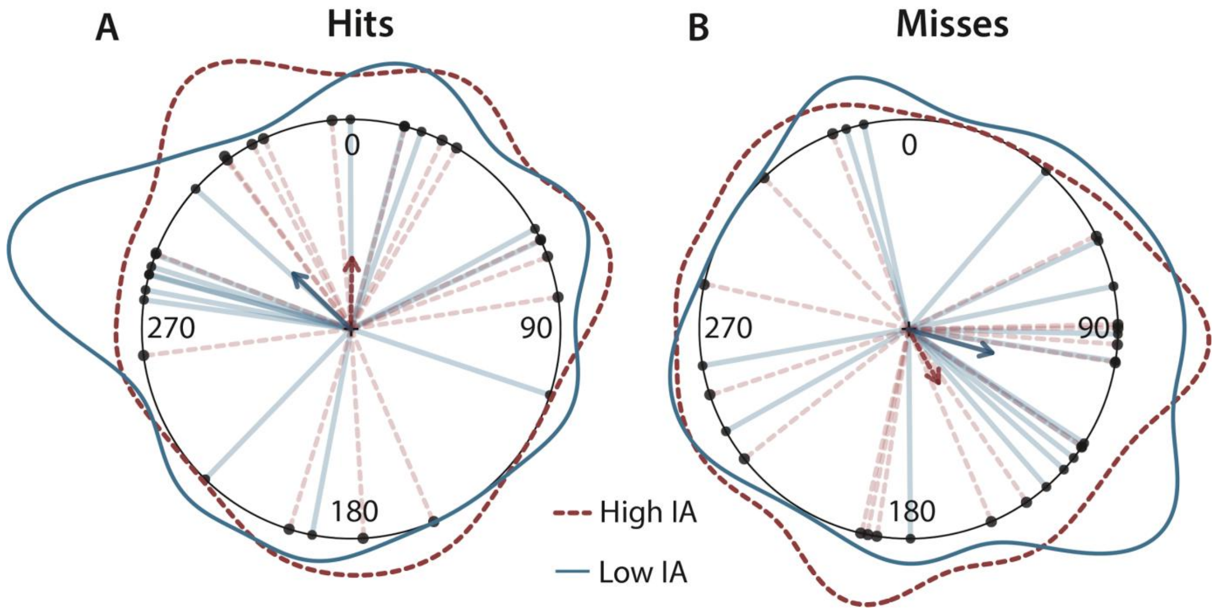


**Figure S1.** Distribution of (**A**) hits and (**B**) misses across the cardiac cycle (i.e, the interval between two R peaks; at 0/360°) separately for subjects with high (dashed red lines) or low (solid blue lines) interoceptive accuracy (IA). Each dot (and line) indicates one participant's mean phase angle. The annular line depicts the distribution of individual means. The darker arrows represent the directions of the group means for hits (high IA: 1°, low IA: 314°) and misses (high IA: 150°, low IA: 105°), with their length indicating the concentration of individual means across the cardiac cycle (hits: high IA – 0.34, low IA – 0.38; misses: high IA – 0.29, low IA – 0.42). Distributions of hits or misses did not differ significantly between the groups with high or low IA.

### 2.3. Differences in hit rates between systole and diastole for high and low interoceptive accuracy groups

A two-way mixed ANOVA of hit rates was conducted, with cardiac phase as a within-subject factor (systole and diastole) and interoceptive accuracy as a between-subject factor (high and low IA). In line with the results reported in the main text of the paper (section 3.3), we found a significant main effect of cardiac phase ($F(1, 30) = 6.36$, $p = 0.017$, $\eta^2_G = 0.006$; Fig. S2) and neither the main effect of IA ($F(1, 30) = 2.50$, $p = 0.960$, $\eta^2_G < 0.001$) nor the interaction between IA and cardiac phase ($F(1, 30) = 1.23$, $p = 0.728$, $\eta^2_G < 0.001$) were significant. These results are consistent with the outcomes of the circular analyses in not showing a significant influence of interoceptive accuracy on variations of conscious somatosensory perception across the cardiac cycle.

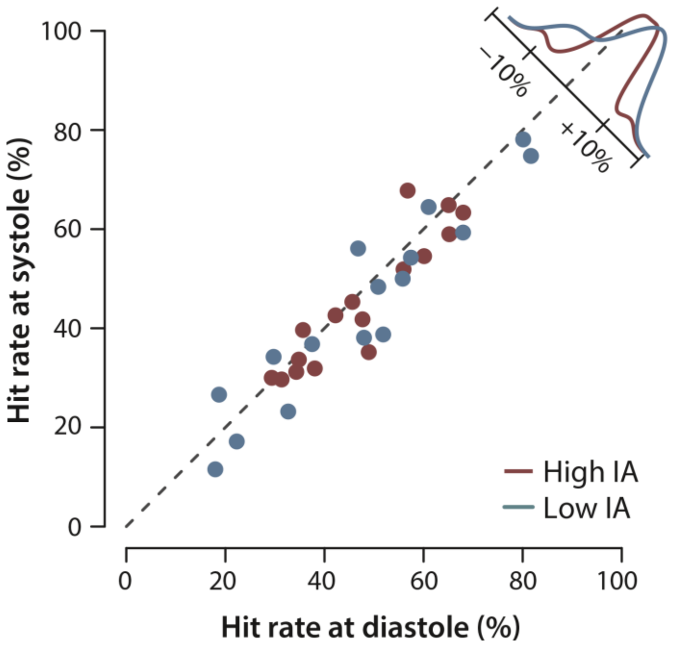


**Figure S2.** Hit rates in diastole and systole for subjects with high (red) or low (blue) interoceptive accuracy (IA). The coordinates of each dot represent a participant's mean hit rate at systole (x-axis) and diastole (y-axis). The dashed lines mark the identity line in hit rate between cardiac phases. The distribution in the upper right corner aggregates the frequencies for high and low IA groups. Hit rates were significantly higher in diastole than in systole but did not differ between the groups with high or low IA.

**REFERENCES CITED IN SUPPLEMENT**

Garfinkel, S.N., Seth, A.K., Barrett, A.B., Suzuki, K., & Critchley, H.D. (2015) Knowing your own heart: distinguishing interoceptive accuracy from interoceptive awareness. *Biological Psychology*, *104*, 65–74. https://doi.org/10.1016/j.biopsycho.2014.11.004

Pewsey, A., Neuhäuser, M., & Ruxton, G.D. (2013). *Circular statistics in R*. Oxford and New York: Oxford University Press.

Schandry, R. (1981) Heart Beat Perception and Emotional Experience. *Psychophysiology*, *18*, 483–488. https://doi.org/10.1111/j.1469-8986.1981.tb02486.x
